# Focal fMRI signal enhancement with implantable inductively coupled detectors

**DOI:** 10.1101/2021.04.02.438254

**Authors:** Yi Chen, Qi Wang, Sangcheon Choi, Hang Zeng, Kengo Takahashi, Chunqi Qian, Xin Yu

**Author notes:** These authors contributed equally to this work (Yi Chen, Qi Wang). Corresponding authors. **Lead corresponding authors:** Dr. Xin Yu, Dr. Chunqi Qian.

## Abstract

Despite extensive efforts to increase the signal-to-noise ratio (SNR) of fMRI images for brain-wide mapping, technical advances of focal brain signal enhancement are lacking, in particular, for animal brain imaging. Emerging studies have combined fMRI with fiber optic-based optogenetics to decipher circuit-specific neuromodulation from meso to macroscales. Acquiring fMRI signal with high spatiotemporal resolution is needed to bridge cross-scale functional dynamics, but SNR of targeted cortical regions is a limiting factor. We have developed a multi-modal fMRI imaging platform with an implanted inductive coil detector. This detector boosts the tSNR of MRI images, showing a 2-3 fold sensitivity gain over conventional coil configuration. In contrast to the cryoprobe or array coils with limited spaces for implanted brain interface, this setup offers a unique advantage to study brain circuit connectivity with optogenetic stimulation and can be further extended to other multi-modal fMRI mapping schemes.

## INTRODUCTION

The combination of fMRI with optogenetics presents a promising tool for mechanistic studies of neuromodulation with cellular and circuit specificity from meso-to-macroscales (Lee, Durand et al. 2010, Ferenczi, Zalocusky et al. 2016, Ryali, Shih et al. 2016, Yu, He et al. 2016, Albers, Schmid et al. 2018, Chen, Sobczak et al. 2019). However, given indirect measurements of neuronal activity with fMRI (Ogawa, Tank et al. 1992, Logothetis, Pauls et al. 2001, Logothetis 2008), it remains ambiguous to represent the circuit-specific brain activation with global hemodynamic patterns, *e.g.* the blood-oxygen-level-dependent (BOLD) functional maps (Logothetis 2010). Besides brainwide functional imaging, high-resolution fMRI of focal brain regions has emerged to map laminar-specific BOLD signal across cortical layers (Silva and Koretsky 2002, Goense and Logothetis 2006, Chen, Wang et al. 2013, Yu, Qian et al. 2014, Huber, Handwerker et al. 2017, Albers, Schmid et al. 2018, Kashyap, Ivanov et al. 2018, Finn, Huber et al. 2019, Sharoh, van Mourik et al. 2019, Yu, Huber et al. 2019, Sangcheon Choi 2021). This unique mapping scheme extends the power of fMRI to show higher specificity when mapping cortical neuronal projection patterns (Yu, Qian et al. 2014, Huber, Handwerker et al. 2017, Sharoh, van Mourik et al. 2019). Recently, there has been an increasing trend to apply laminar fMRI for human brain mapping with either top-down or bottem-up tasks (Huber, Handwerker et al. 2017, Kashyap, Ivanov et al. 2018, Finn, Huber et al. 2019, Sharoh, van Mourik et al. 2019, Yu, Huber et al. 2019). To elucidate the neural circuit regulation of layer-specific BOLD responses, it is crucial to implement the optogenetic laminar fMRI in animal models.

An ongoing challenge of laminar fMRI is the limited signal-to-noise ratio (SNR) when sampling the layer-specific BOLD signal with high resolution. With the high field (>11.7 Tesla) MRI, a line-scanning fMRI method has been developed to extract BOLD signal with sufficient SNR along cortical layers of designated cortices (Yu, Qian et al. 2014, Sangcheon Choi 2021). Albers et al has applied the line-scanning fMRI to detect the layer-specific BOLD responses driven by optogenetic stimulation in rodent brains (Albers, Schmid et al. 2018). However, given the implantation of optical fiber, it remains challenging to apply the advanced cryoprobe (Darrasse and Ginefri 2003, Baltes, Radzwill et al. 2009) or multi-array radio frequency (RF) coils (Roemer, Edelstein et al. 1990, Gareis, Wichmann et al. 2007) for focal signal enhancement. Also, even with the surface coil designed to accommodate optical fiber implantation (Yu, He et al. 2016, Chen, Sobczak et al. 2019, Chen, Sobczak et al. 2020), the adhesive surgical material used to fix the fiber at the skull creates a large distance between the surface coil and brain regions, which further restricts the acquisition of laminar fMRI signal with sufficient SNR.

It is well established that inductively coupled detectors (ICD) can be placed near the targeted region of interests (ROI) to relay locally detected MR signals to the external surface RF coil wirelessly, thus reducing hardware complexity and boosting sensitivity (Schnall, Barlow et al. 1986, Wirth, Mareci et al. 1993, Volland, Mareci et al. 2010, Ginefri, Rubin et al. 2012, Mett, Sidabras et al. 2016). Examples inside high field MRI scanners include an inductively coupled MR coil system for imaging and spectroscopic analysis of an implantable bioartificial construct in mice peritoneal cavity at 11.1 T (Volland, Mareci et al. 2010), an inductively coupled coil pair to obtain magnetic resonance phantom images.at 9.4 T (Mett, Sidabras et al. 2016), and an inductively-coupled coil at the level of the interhemispheric cleft to obtain rat brain anatomical images at 7 T (Ginefri, Rubin et al. 2012). All these studies demonstrated sensitivity enhancement in very close proximity to the targeted ROI with the inductive coils. Despite of its promising prospects in other MRI research, to the best of our knowledge, it has not been utilized on a multi-modal platform to offer a novel and elegant solution to the challenges mentioned above.

Hence, as a proof-of-concept demonstration, to improve MR detection sensitivity for cross-scale brain mapping, we presented an optimized multi-modal fMRI platform with ICD. For verification of its feasibility and SNR improvement, whole brain echo-planar imaging (EPI) and line-scanning fMRI were performed in anesthetized rat brains. Moreover, as an *in vivo* benchmark application of ICD against conventional surface coil, we have embedded the ICD with optical fiber implantation to show a 3-fold enhancement of SNR in optogenetically-driven laminar BOLD fMRI experiments.

## MATERIAL AND METHODS

### Flexible implanted inductive coil design and fabrication

The inductive coil design we used is based on wireless inductive coupling theorem, with smaller-dimention circuit to maintain detection sensitivity (**Fig.1a**). The inductive coil was tuned to 599.58 MHz by the formula: 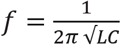, matching the resonance frequency delivered by a transceiver coil in 14.1 T scanner. Hence, the dimension of conductor was determined by value of inductance *L*) and the tuning capacitance was adjusted to maintain constant resonance frequency. The fabrication process is as below: the conductor was initially printed with copper material on PCB (by Electronics Workshop of Max Planck Institute for Biological Cybernetics, Tübingen, Germany), in circle with a dimension of 1 mm strip width, 6 mm outer diameter; the capacitor was implemented by an unbiased diode (BBY5302VH6327XTSA1, Infineon Technologies, Germany); all assembly process of the designed circuit (**Fig.1b**) is done by soldering. For the expression of on-bench performance of inductive coil, its frequency response profile was colormapped under different distance separations from 0-25 mm with a stepsize of 1 mm, using *imagesc* function of MATLAB (**Fig. 1c**).

### Animals

The study was performed in accordance with the German Animal Welfare Act (TierSchG) and Animal Welfare Laboratory Animal Ordinance (TierSchVersV). This is in full compliance with the guidelines of the EU Directive on the protection of animals used for scientific purposes (2010/63/EU). The study was reviewed by the ethics commission (§15 TierSchG) and approved by the state authority (Regierungspräsidium, Tübingen, Baden-Württemberg, Germany). A 12-12 hour on/off lighting cycle was maintained to assure undisturbed circadian rhythm. Food and water were obtainable ad libitum. A total of 6 male Sprague–Dawley rats acquired from Charles River were used in this study.

### Viral injection and immunohistochemistry

AAV5.CaMKII.hChR2 (H134R)-mCherry (100 μL at titer ≥ 1 × 10 ^13^ vg/mL) from Addgene was injected in 4-week-old rats intracerebrally in the right somatosensory forepaw region. Rats were anesthetized with 1.5-2% isoflurane via nose cone and placed on a stereotaxic frame, an incision was made on the scalp to expose the skull. Craniotomies were performed with a pneumatic drill so as to introduce minimal damage to cortical tissue. A volume of 0.6-0.9 μL was injected using a 10-μL syringe and 33-gauge needle to the stereotaxic coordinates: 0 mm posterior to Bregma, 3.8-4.0 mm lateral to the midline, 0.8-1.4 mm below the cortical surface. After injection, the needle was left in place for approximately 5 mins before being slowly withdrawn. The craniotomies were sealed with bone wax and the skin around the wound was sutured. Rats were subcutaneously injected with antibiotic and painkiller for 3 consecutive days to prevent bacterial infections and relieve postoperative pain.

To verify the phenotype of the transfected cells, opsin localization and optical fiber placement, perfused rat brains were fixed overnight in 4% paraformaldehyde and then equilibrated in 15% and 30% sucrose in 0.1 M PBS at 4°C. 30 μm-thick coronal sections were cut on a cryotome (CM3050S, Leica, Germany). Free-floating sections were washed in PBS, mounted on microscope slides, and incubated with DAPI (VectaShield, Vector Laboratories, USA) for 30 mins at room temperature. Wide-field fluorescent images were acquired using a microscope (Zeiss, Germany) for assessment of ChR2 expression in FP-S1. Digital images were minimally processed using ImageJ to enhance brightness and contrast for visualization purposes.

### Animal preparation, inductive coil/fiber optic implantation for fMRI

Anesthesia was first induced in the animal with 5% isoflurane in the chamber. The anesthetized rat was intubated using a tracheal tube and a mechanical ventilator (SAR-830, CWE, USA) was used to ventilate animals throughout the whole experiment. Femoral arterial and venous catheterization was performed with polyethylene tubing for blood sampling, drug administration, and constant blood pressure measurements. After the surgery, isoflurane was switched off, and a bolus of the anesthetic alpha-chloralose (80 mg/kg) was infused intravenously. A mixture of alpha-chloralose (26.5 mg/kg/h) and pancuronium (2 mg/kg/h) was constantly infused to maintain anesthesia/keep the animal anesthetized and reduce motion artifacts.

Before the animal was transferred to the MRI scanner, it was placed on a stereotaxic frame and the scalp was opened. For eletrcial stimulation in **Fig.2**, the 6-mm single loop inductive coil was directly placed on the right FP-S1 region and secured with adhesive gel (**Fig. 1a**) beneath the 22-mm surface coil (**Fig.1a,b**). For optogenetic experiment in **Fig. 3**, one ~1.5-mm diameter burr holes was drilled on the skull and the dura was carefully removed. An optical fiber with 200-μm core diameter (FT200EMT, Thorlabs, Germany) was inserted into the FP-S1 through the hole of the 6-mm inductive coil, at coordinates of 0 mm posterior to Bregma, 4 mm lateral to the midline and 1.2-1.4 mm below the cortical surface. An adhesive gel was used to secure the fiber to the skull with the inductive coil beneath. While the other 6-mm inductive voil was placed on the projected FP-S1 in the left hemisphere. The eyes of the rats were covered to prevent stimulation of the visual system during the optogenetic fMRI.

### MRI acquisition

All images were acquired with a 14.1 T/26 cm horizontal bore magnet interfaced to an Avance III console and equipped with a 12-cm gradient set capable of providing 100 G/cm over a time of 150 μs. A transceiver single-loop surface coil and inductive coils as described above were used to acquire MRI images.

The FLASH images in **Fig. 1d** for phantoms were acquired with the following parameters: TR/TE 100/5 ms, excitation pulse angle 30°, slice thickness 0.2 mm, FOV 4 cm × 4 cm and matrix 256 × 256. Functional images in **Fig. 2d, e** were acquired with a 3D gradient-echo EPI sequence with the following parameters: Echo Time 11.5 ms, repetition time 1.5 s, FOV 1.92 cm × 1.92 cm × 1.92 cm, matrix size 48 × 48 × 48, spatial resolution 0.4 mm × 0.4 mm × 0.4 mm. For anatomical reference, the RARE sequence was applied to acquire 48 coronal slices with the same geometry as that of the fMRI images. The FLASH-based bilateral line-scanning method was combining 2 saturation RF pulses to dampen the MR signal outside the regions of interest (**Fig. 2e**) with the following parameters: TR/TE 100/5 ms, excitation pulse angle 30°, slice thickness 1 mm, FOV 6.4 mm × 3.2 mm and matrix 64 × 32. The phase-encoding gradient was turned off. Functional activation was detected by performing electrical stimulation on left forepaw (3 Hz, 4 s, 300 μs width, 2.5 mA) and optogenetic stimulation on right FP-S1 (2 Hz, 6 s, light pulse width 10 ms, 30 mW) to activate the neurons expressing AAV.CaMKII.ChR2.mCherry (**Fig.3a**). BLOCK design was 1 second pre-stimulation, 4 (6 for optogenetic stimulation) second stimulation and 15 (13 for optogenetic stimulation) second post-stimulation, *i.e.,* 20 s for each epoch and 32 epochs for the whole trial (10 m 40 s).

### Data analysis

All signal processing and analyses were implemented in Matlab MATLAB software (Mathworks, Natick, MA) and Functional NeuroImages software (AFNI, NIH, USA). For evoked fMRI analysis for **Fig. 2d,e** and **Fig. S1**, the hemodynamic response function (HRF) used was the default of the block function of the linear program 3dDeconvolve in AFNI. BLOCK (L, 1) computes a convolution of a square wave of duration L and makes a peak amplitude of block response = 1, with g(t) =*g*(*t*) = *t^q^e^-t^*/[*q^q^e^-q^*], where q = 4. Each beta weight represents the peak height of the corresponding BLOCK curve. The HRF model is defined as follows:

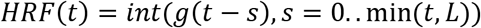

For layer-specific BOLD analysis, the boundaries of cortex were defined at the reference positions of both cortical surface and corpus callosum, as the cortical surface was determined at half maximum signal intensity of intensity profile of fMRI (Yu, Qian et al. 2014), rectified by 0.2 mm offset from corpus callosum center (half width of corpus callosum). As a function of time, fMRI percentage dynamics were calculated based on the following equation: *SI_percetage_* = (*SI_baseline_*)/*SI_baseline_*, Where *SI_i_* denotes signal intensity at the *i^th^* time point, *i* = 0:0.1:19 s for each epoch, and *SI_baseune_* indicates the mean value of signal intensity at pre-stimulation duration (1s). Such time series of activation were presented after voxel-wise dynamics of the entire cortex being averaged and color-coded (**Fig. 2f** top right, and **Fig. 3b** top), through whole scan duration (**Fig. 2f** top left, **Fig. 3a** top left and right). Likewise, laminar-specific activation features were averaged within each layer correspondingly (**Fig. 3c**). For epoch-wise spatiotemporal activations, time series were averaged with respect to each voxel and were color-mapped (*imagesc* function in MATLAB), where signal intensity was expressed in percentages (**Fig. 2f** bottom right and **Fig. 3b** bottom). Due to the contribution of vascular partial volume effect, higher values of percentage changes exist in voxels next to cortical surface.

For resting-state fluctuation of hemodynamics, z-normalized time series (*zscore* function in MATLAB) of a single trial were demeaned to present overall cortical fluctuation (**Fig. 2g** top). As to the spatiotemporal map with whole scan duration, bandpass filter of 0.01-0.1 Hz was applied through zero-phase digital filtering (*filtfilt* function in MATLAB), with intensity values being standardized between 0 and 1 per voxel of cortex (**Fig. 2f** lower left, **Fig. 3a** lower left and right).

Comparison of SNR in **Fig. 2c** and **Fig. 1d** was illustrated by bar plot containing the enhanced region and the normal one encircled by identical geometries. SNR was calculated by:

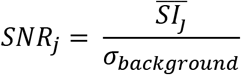

where 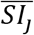 stands for mean of signal intensity in the regions of interest labeled by *j*, indicating enhanced or uncovered region, and *σ_background_* for standard deviation of signal intensity in a sample of background; RF pulse attenuation was automatically adjusted by console of BRUKER system. SNR values of individual animals were plotted as black dots, and error bars were made by stdandard deviation (s.d.) of all animals.

Comparison of temporal Signal-to-Noise-Ratio (tSNR) in **Fig. 2g** (right) and **Fig. 3d** was made between before and after inductive coupling enhancement, which was calculated as:

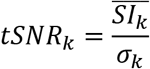

Where 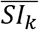 stands for the averaged amplitude of signal time series during the scan of corresponding animal, and *σ_k_* for standard deviation of corresponding time series, both variants were made after averaging the signal intensity over the entire cortical depth.

Statistical significance was calculated by applying paired-sample t-test for both sets of data (*ttest2* function in MATLAB), where shading region covers a range of paired voxels with distinct levels of statistical significance (**Fig. 2g** right and **Fig. 3 d**), *i.e.,* **P < 0.01. Likewise, bar markers of significance level was calculated after averaging tSNR value of animals in each group, with errobars expressing s.d. of individual trials.

## RESULTS

### Design, characterization and *ex vivo* evaluation of the inductive coil detector

We fabricated a 6-mm single-ring ICD whose performance under smaple loading conditions was evaluated both on bench and inside a 14.1-T scanner. **Fig. 1a** demonstrates the proposed single-ring ICD and conceptual diagram for bench measurements. The proposed setup consists of a single-ring ICD and a conventional tranceiving surface coil, while NMR signals from the insulated ICD can be transmitted to the external coil through mutual inductive coupling (**Fig. 1b**). The single-loop ICD was placed on top of a 1% agarose gel phantom while the conventional surface coil above the phantom with a certain distance separation would provide RF excitation pulse and receive amplified signals from ICD (**Fig. 1a-b**). No modifications to the scanner interface were required. As shown in **Fig. 1c**, the strong coupling effect between the ICD and surface coil could be maintained for a distance as large as 12 mm. A representative image for the gel phantom is shown in **Fig. 1d** (left), and the relative SNR from the focal region below the ICD is shown in **Fig. 1d** (right). For distance separations beyond 3 mm, attenuation of RF excitation efficiency is compensated by increased excitation power, maintaining almost constant SNR for a range of distance separations. The *in vitro* test result detects the effective range of distance between the ICD and the surface coil (<12 mm) for inductive coupling.

**Fig.1.**
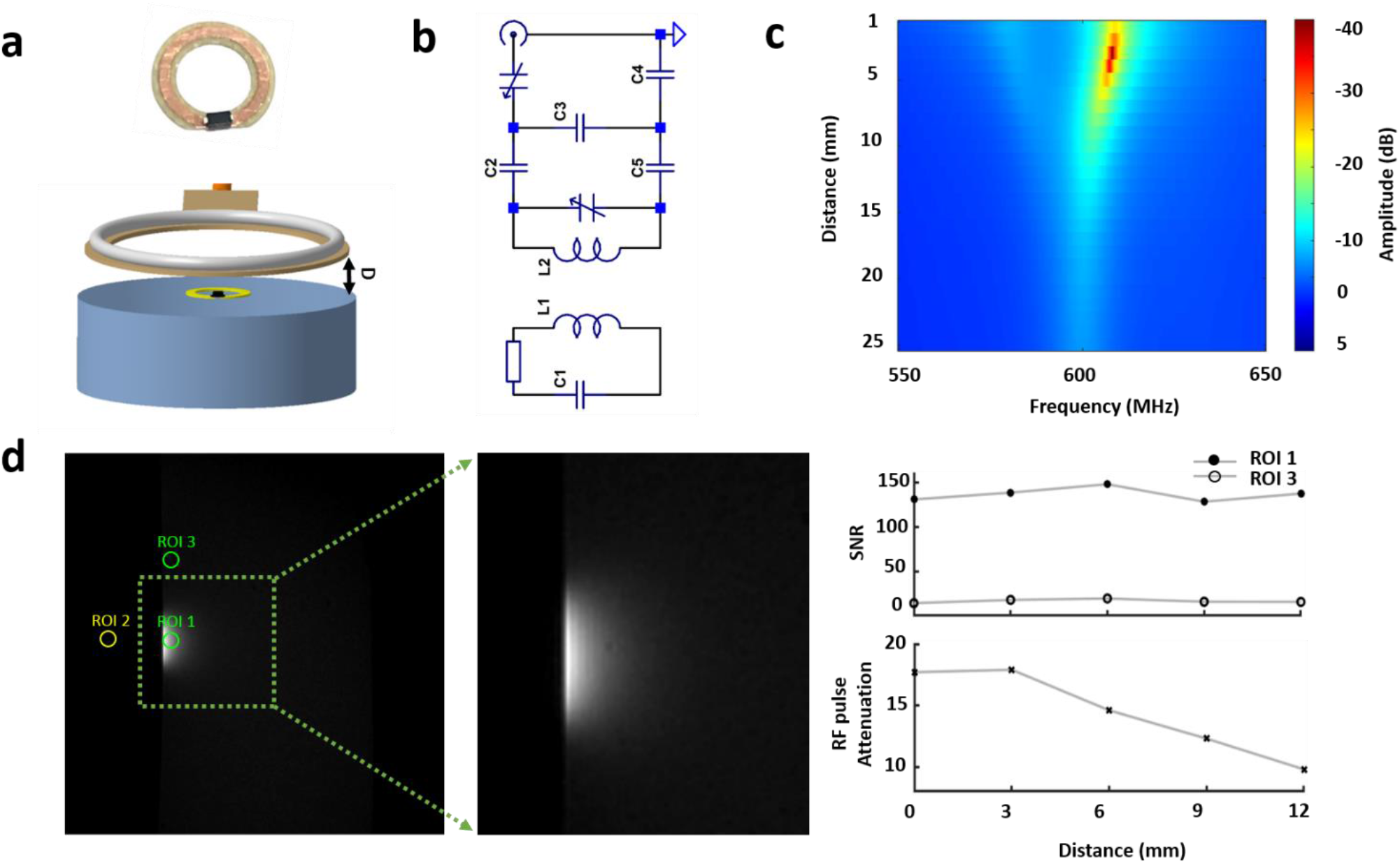
Evaluation of inductive coil under different distance separations. (a) Photograph of IC (upper) and schematic drawing of the phantom test arrangement with the ICD placed on the surface of a 1% agarose gel. (b) Diagrams of IC implemented as a single resonant circuit coupled inductively to the external RF surface coil. (c) Coupling performance varies with the increased distant separations between the inductive loop and the transceiver coil. (d) Intensity of FLASH 2D image acquired from the gel phantom shows significantly higher SNR inside enhanced region (ROI 1) than the unenhanced region (ROI 3). The ICD was placed on the gel while the surface coil was separated from the gel surface by 3 mm. Left: phantom image with regions of interest for performance comparison. Right upper, spatial SNR comparison between regions when the two coils are separated by increasingly large distances. Bottom, the required attenuation of RF pulse power changes as a function of distance separation.

### *In vivo* evaluation of inductive coil detector with layer-specific BOLD mapping in rat brain

Next, ICD was evaluated *in vivo* to measure enhanced fMRI signal in the right forepaw somatosensory cortex (FP-S1) with unilateral electrical forepaw stimulation and during rest in anesthetized rat brains (**Fig.2**). A 6-mm single loop inductive coil was embedded beneath the 22-mm surface coil at the right FP-S1 (**Fig.2a**). By relaying locally detected MR signals to the external surface coil, the focal intensity enhancement by this 6-mm diameter ICD was detected throughout the 2 mm cortex from anatomical FLASH MR images (**Fig. 2b)**. The focal signal intensity in ROI 1 below the ICD in the right FP-S1 was siginificantly higher than that of ROI 2 in the left hemisphere without ICD (**Fig. 2b, c**). As shown in **Fig.2d** (**Fig. S1** for another 2 rats), the anatomical image with superimposed BOLD functional maps demonstrated that the ICD-enhanced region was well overlapped with the most activated FP-S1. Besides the EPI-based BOLD, we acquired laminar-specific BOLD responses with 100-ms temporal resolution and 100-μm spatial resolution using line-scanning fMRI (**Fig.2e**). The time courses and line profile-based 2D fMRI maps across cortical layers were presented in stimulation (**Fig 2f**) and resting state conditions (**Fig 2g**). The tSNR for resting state brain fluctuation in the cortex illustrates the ICD is up to 2-fold more sensitive than the surface coil only condition (**Fig.2g** and **Fig. S2**).

**Fig.2.**
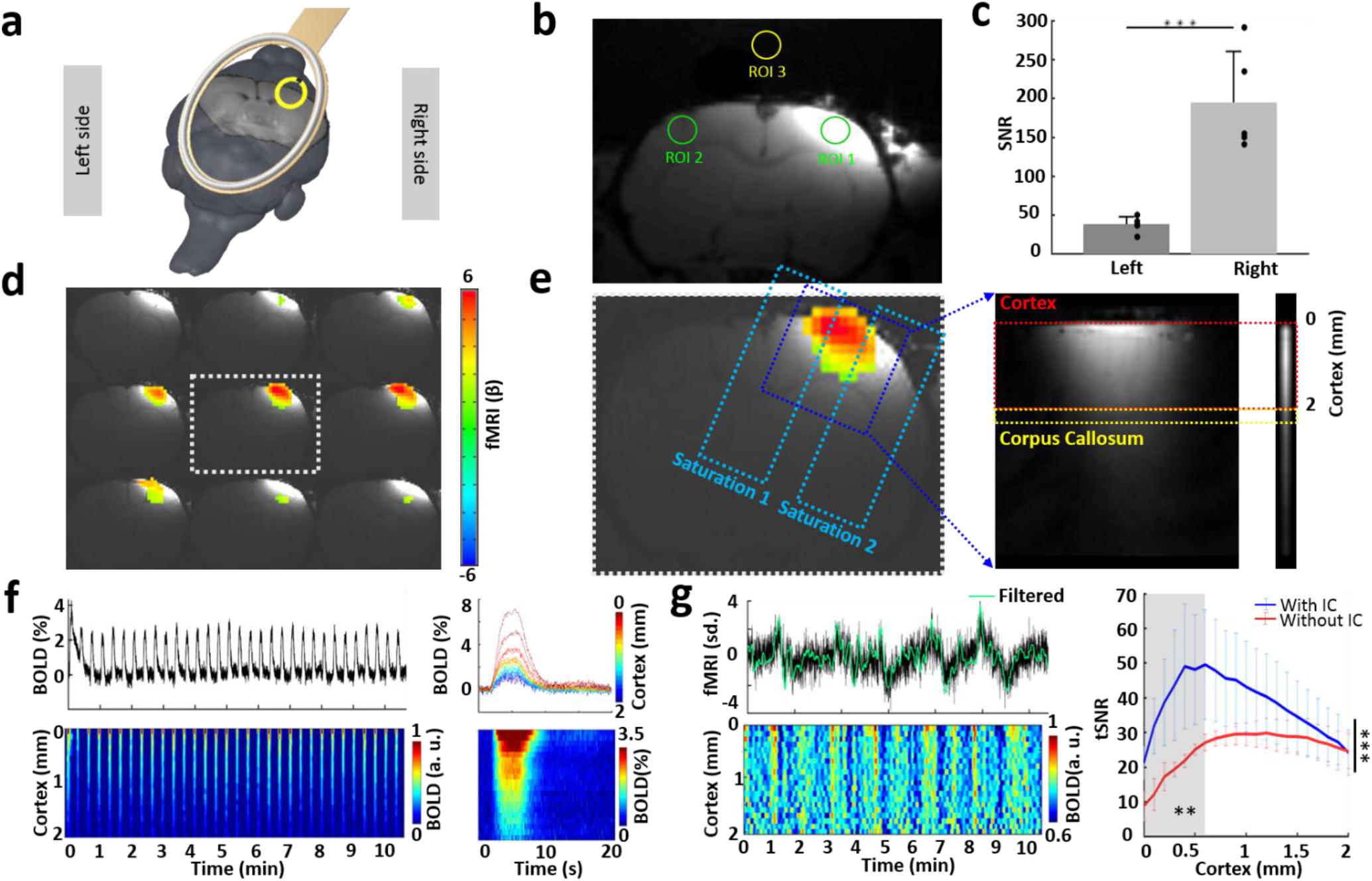
BOLD responses detected using unilateral line-scanning method in enhanced region. (a) Inductive coil covers FP-S1 region on the right hemisphere with a surface coil covering the whole rat brain. (b) A representative anatomic image shows enhanced focal intensity in the right FP-S1 region (ROI 1). (c) Significantly higher SNR in ROI 1 with inductive coil (right) than in ROI 2 (left) (Paired-sample t-test, ***P = 7 × 10^-4^, n = 5 rats, mean ± SD). (d) The color-coded BOLD activation map with left forepaw electrical stimulation (3 Hz, 4s, 2 mA) superimposed on the anatomical images (FLASH). (e) The procedure to set up the unilateral line-scanning method. (f) Averaged BOLD percentage change from all 20 voxels in the cortex for 32 epochs (top left) and for each epoch along the cortical depth (top right, 20 voxels, 2 mm). Normalized spatiotemporal map (bottom left) along cortical depth for the trial (bottom left) and the epoch (bottom right) (n = 3 rats, 17 trials, 32 epochs, 10min 40s per trial). (g) Averaged time course (top left) extracted from right cortex shows the hemodynamic fluctuation in the absence of stimulation and its voxel-specific normalized spatiotemporal map (bottom left) throughout the cortex (20 voxels, 2 mm). Right, the tSNR with inductive coil (blue) is significantly higher than our previous results acquired with surface coil (red). It is noteworthy that SNR close to the corpus callosum is compatible (paired-sample t-test, ***P <0.001, **P < 0.01, n = 3 rats, mean ± SD).

### Evaluation of ICD at multi-modal fMRI platform with optogenetic-driven layer-specific BOLD mapping

The high sensitivity of ICD was verified by combining optical fiber-based optogenetic stimulation and bilateral line-scanning method (Sangcheon Choi 2021). We embedded two ICD to cover the FP-S1 of both hemispheres with optical fiber implanted to target the right FP-S1 expressing channelorhodopsin-2 (ChR2) (**Fig. 3a**, middle). Upon optogenetic stimulation, robust BOLD responses were detected in the right FP-S1 region (**Fig. 3a**, right), as well as evoked BOLD signal in the left FP-S1 (**Fig. 3a**, left). The averaged line profile-based 2D fMRI maps (**Fig. 3b**) and time courses (**Fig. 3c**) were displayed to repsent the layer-specific hemodynamic responses. Interestingly, the evoked BOLD signal in the left FP-S1 ipsilateral to the stimulated forepaw showed salient post-stimulus undershoots in L2/3 and L5, indicating a transcallosal projection-mediated interhemispheric inhibition (Karayannis, Huerta-Ocampo et al. 2007, Palmer, Schulz et al. 2012, Chen, Sobczak et al. 2020). Also, the laminar-specific tSNR with ICD showed up to 3-fold SNR increase over the bilateral line-scanning fMRI signals detected only with surface coil (**Fig. 3d**). This result demonstrates the unique advantage to boost the SNR with ICD-based optogenetic laminar-fMRI.

**Fig.3.**
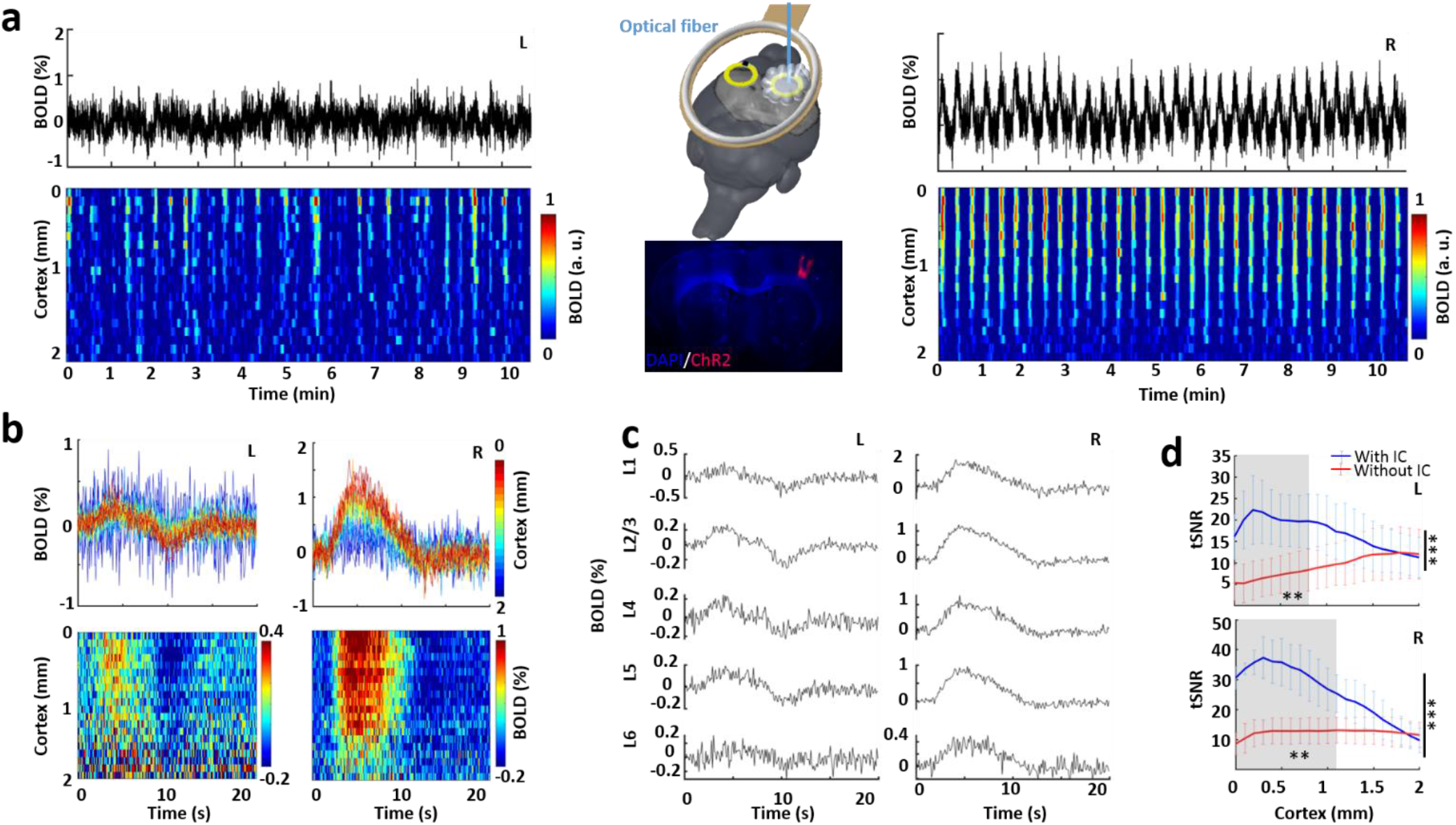
Optogenetically evoked BOLD responses in bilateral enhanced FP-S1 regions. (a) Averaged time course (top) and normalized spatiotemporal map (bottom) of bilateral BOLD responses in FP-S1 regions induced directly by optogenetic stimulation in the right hemisphere (right) and projected left hemisphere (left, n = 3 rats, 49 trials, 32 epochs, 2 Hz, 6 s, 10 ms light pulse, 30 mW). Middle upper, schematic drawing of two inductive coils stuck adjacent to bilateral hemispheres, with optical fiber (blue) inserted to the right FP-S1 region. Middle lower, a representative wide-field fluorescence image illustrates robust ChR2-mCherry expression in the right FP-S1. (b) fMRI percentage-change time courses (top) and maps (bottom) for epoch in each voxel along cortical depth on both hemispheres (n = 3 rats, 49 trials). (c) Averaged epoch-wise time courses showing the different laminar-specific responses of both hemispheres (n = 3 rats). (d) The comparison for both hemispheres shows significantly higher tSNR with (blue) than without (red) implanted inductive coils (paired-sample t-test, ***P <0.001, **P < 0.01, n = 3 rats, mean ± SD).

## DISCUSSION

In this study, we have presented a multi-modal platform by combining optogenetics and fMRI in animals with implanted ICD (Inductively Coupled Detector) to increase the SNR. Unlike the global functional mapping scheme, laminar fMRI has been developed to extract the layer-specific BOLD signal from focal brain regions with high spatial resolution (Silva and Koretsky 2002, Goense and Logothetis 2006, Chen, Wang et al. 2013, Yu, Qian et al. 2014, Huber, Handwerker et al. 2017, Albers, Schmid et al. 2018, Kashyap, Ivanov et al. 2018, Finn, Huber et al. 2019, Sharoh, van Mourik et al. 2019, Yu, Huber et al. 2019). Conventional surface coils have enabled the focal signal enhancement, in particular, with miniaturied design to be positioned close to cortical regions (Ackerman, Grove et al. 1980, Chen, Sobczak et al. 2019). Also, surface coils can accommodate the optogenetic setup with fiber bundles passing through the open area of the coil, which is not compatible with the insulated ceramic holder that is required for cryo-probe to maintain its superconducting feature inside cyrogen (Peter Styles 1989). It should also be noted that when optical fiber was implanted to target brain region, a large distance separation between the RF coil and cortical regions will be introduced, leading to significantly reduced sensitivity of surface coil for focal brain signal detection. The ICD was developed to overcome this limitation through implantation in close proximity to the cortical region, which can be wirelessly coupled to an external surface coil (Wirth, Mareci et al. 1993, Volland, Mareci et al. 2010, Ginefri, Rubin et al. 2012, Mett, Sidabras et al. 2016). This ICD-based mapping scheme can also be extended to target multiple brain regions to enable cross-scale brain dynamic mapping with a simple experiment setup (**Fig. 3**), showing 2-3 fold sensitivity gain over conventional surface coils (**Fig. 2d** and **Fig. 3d**). Also, ICD implantation with the optical fiber simplifies the fMRI imaging setup and frees open space for other complementary imaging modalities, *e.g.,* fluorescent calcium recording (Schulz, Sydekum et al. 2012, Albers, Wachsmuth et al. 2018, Wang, He et al. 2018, Lake, Ge et al. 2020) and pupillometry (Pais-Roldan, Takahashi et al. 2020). Thus, we estliabhed an *in vivo* benchmark ICD application for multi-modal high-resolution fMRI studies.

To validate the unique advantage of ICD-based focal signal enhancement, we implemented bilateral line-scanning fMRI with optogenetics. To date, the spatiotemporal dynamic patterns of evoked fMRI signal through transcallosal projections have not been fully investigated, in particular, the layer-specific fMRI signal based on the regulation of interhemispheric excitatory/inhibitory balance. Electrophysiology in brain slices have elucidated that CC-mediated glutamatergic excitatory postsynaptic potentials are followed by elongated GABA-mediated inhibitory postsynaptic potentials (Kawaguchi 1992, Kumar and Huguenard 2001, Karayannis, Huerta-Ocampo et al. 2007, Palmer, Schulz et al. 2012). The optogenetically driven callosal activity has been used to disentangle circuit-specific interhemispheric inhibitory effects, *e.g.*, in the auditory cortex (Rock and Apicella 2015), prefrontal cortex (Lee, Gee et al. 2014) and hindlimb somatosensory cortex (Palmer, Schulz et al. 2012). Interestingly, Iordanova et al. revealed transcallosal-mediated hemodynamic responses into three major categories: negative, biphasic, and no-response, with hemoglobin-based optical intrinsic signal imaging following optogenetic stimulation (Iordanova, Vazquez et al. 2018). Also, Hoffmeyer and colleagues reported the nonlinear neurovascular coupling in rat sensory cortex upon direct electrical stimulation of transcallosal pathways (Hoffmeyer, Enager et al. 2007). Previously, we reported the frequency-dependent BOLD fMRI responses through direct optogenetic stimulation of the callosal projection fibers with whole brain EPI-fMRI. Here, the ICD-based laminar fMRI enables the detection of layer-specific BOLD signal mediated by transcallosal projection (**Fig. 3**). The salient post-stimulus undershoots detected in Layer 2/3 and 5 present the unique coupling feature of hemodynamic responses with interhemispheric excitatory/inhibitory balance (Chen, Sobczak et al. 2020).

Several limitations pertaining to the initial use of ICD in multi-modal platform should be considered when interpreting the results of this work and for future optimization of the ICD in high field MRI scanners. Firstly, although the ICD has 2-3 fold sensitivity gain over external surface coil, its coupling with the external surface coil is passive, thus limiting its effective distance separation from the external surface coil. On the other hand, it is well established that Wirelessly Amplified NMR Detector (WAND) can efficiently amplify MRI signals, leading to an additional >3-fold sensitivity gain over passive coupling (Qian, Yu et al. 2013, Qian, Yu et al. 2020) especially for larger distance separations. Therefore, it will be helpful to utilize the WAND to improve the effective operation range of implantable detectors. Secondly, even for passive coupling, there is still room for further improvement. For example, the angle between the external surface coil and the inductive coil should be as parallel as possible, ensuring higher coupling coefficicent between the two coils. Normally, the parallel arrangement can be implemented readily in phantoms. For *in vivo* experiment, however, the inductive coil is implanted to attach the curved skull, making parallel arrangement harder. This explains the 6-7 fold sensitivity gain for *in vitro* experiments (**Fig. 1d**) in comparison to only 2-3 fold enhancement for *in vivo* experiments (**Fig. 2g** and **Fig. 3d**). Moreover, though the averaged tSNR was increased, because the different coupling coefficient for each in vivo experiments, the SEM with IC has larger variance across the cortical depth that SEM without IC (**Fig. 2g** and **Fig. 3d**). Thirdly, although we only discussed focal signal enhancement with ICD, it is also possible to develop the wireless, implantable RF coil array to enhance detection sensitivity of the entire brain (Bulumulla, Fiveland et al. 2015).

## CONCLUSIONS

In summary, we present the multi-modal imaging scheme with implanted ICDs, yielding high-resolution structural and functional images of the rat brain with high SNR. The ICD-based mapping scheme facilitate optogenetic-driven brain connectivity studies with simplified environmental setup, which can also be used to combine fMRI with real-time microinjection, fiber optic-mediated optical imaging, pupillometry, and electrophysiological recordings to decipher brain function across different scales.

## Data availability

The data that support the findings of this study are available from the corresponding authors upon request.

## Code availability

The related image processing codes are available from the corresponding author upon request.

## Author Contributions

X.Y. and C.Q. designed and supervised the research, Y.C. and Q.W. performed animal experiments, acquired data, analyzed data, S.C., H.Z., and K.T. provided conceptual and key technical support, X.Y., C.Q., Y.C. and Q.W. wrote the manuscript.

## ACKNOWLEDGEMENTS

This research was supported by NIH Brain Initiative funding (RF1NS113278-01, R01 MH111438-01), and the S10 instrument grant (S10 MH124733-01) to Martinos Center, German Research Foundation (DFG) Yu215/3-1, Yu315/2-1, BMBF 01GQ1702, and the internal funding from Max Planck Society. This project has received funding from the European Union Framework Programme for Research and Innovation Horizon 2020 (2014-2020) under the Marie Skłodowska-Curie Grant Agreement No.896245. We thank Dr. MH. Frosz, Mr. J Walzog, Dr. R. Pohmann, Dr. J. Engelmann, Dr. N. Avdievitch and Ms. H. Schulz for technical support, Dr. P. Douay and Ms. R. König for animal support, the AFNI team for the software support.

## Supplementary Information

**Figure S1.**
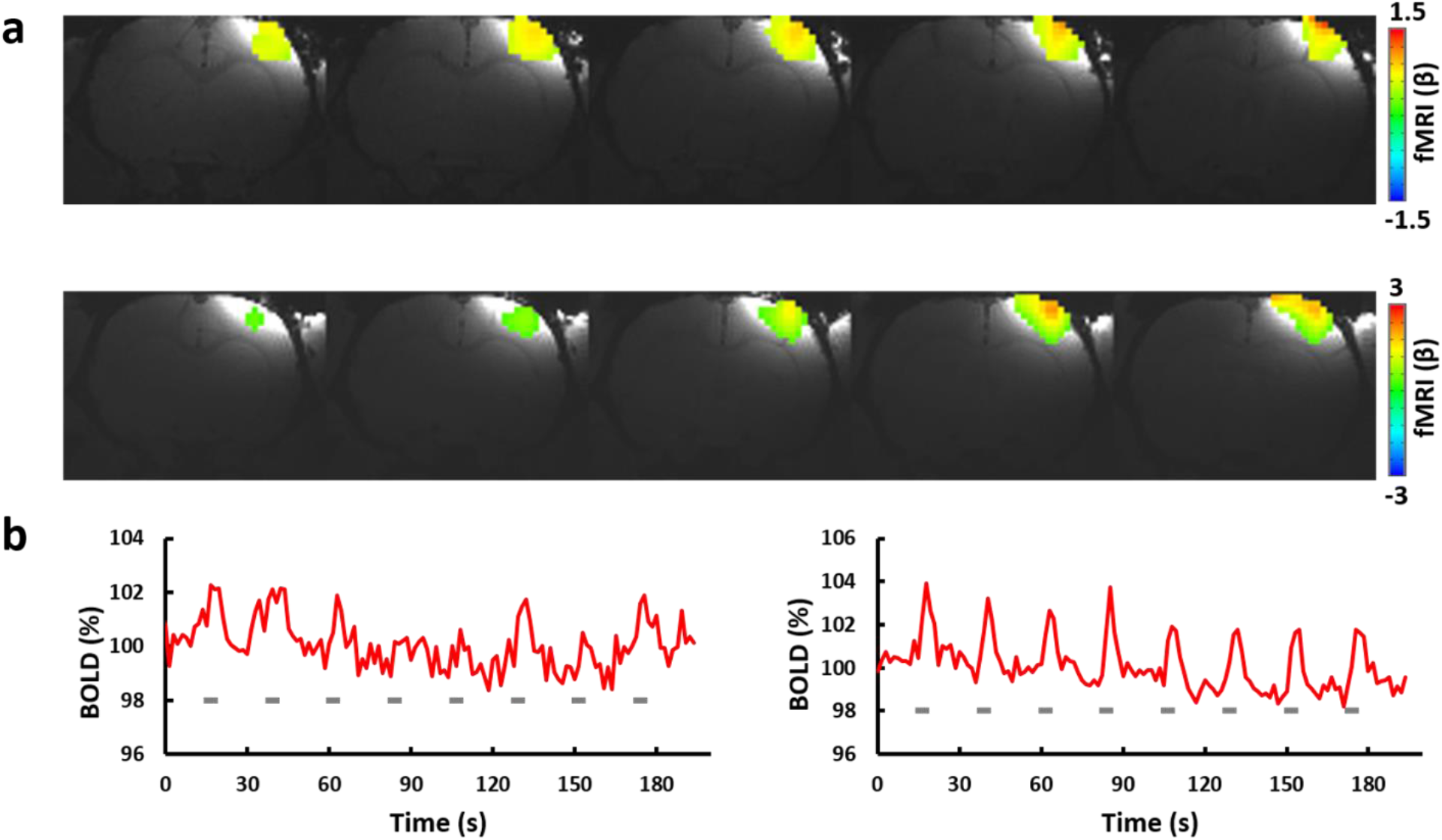
Verification of ICD in another two rats. (**a**) BOLD map upon electrical stimulation on the left forepaw. The anatomical image with superimposed whole brain EPI demonstrates the signal enhanced region that was well matched to the most activated region. (**b**) Averaged BOLD percentage change from the cortex for 8 epochs for rat #1 (left) and rat #2 (right). Grey bars indicate the sensory stimulation.

**Figure S2.**
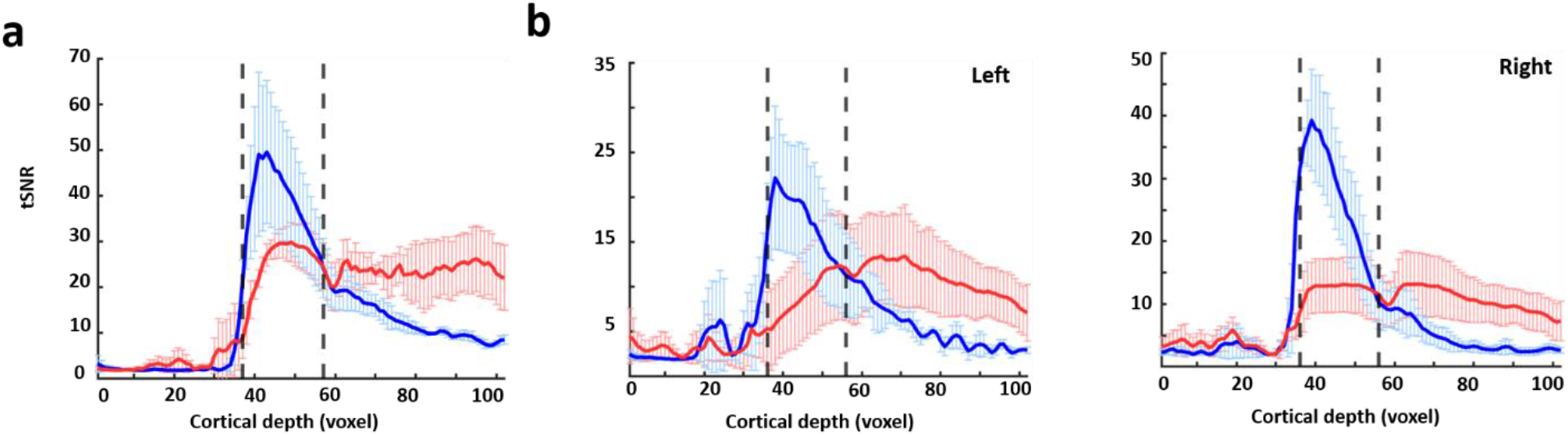
The tSNR with inductive coil (blue) is significantly higher than our previous results acquired with surface coil (red) in the cortex, but not in the subcortical brain regions. (**a**) tSNR comparison for sensory evoked brain activity in the right FP-S1. (**b**) tSNR comparison for optogenetically evoked brain activity in FP-S1 in both hemispheres.

**Figure S3.**
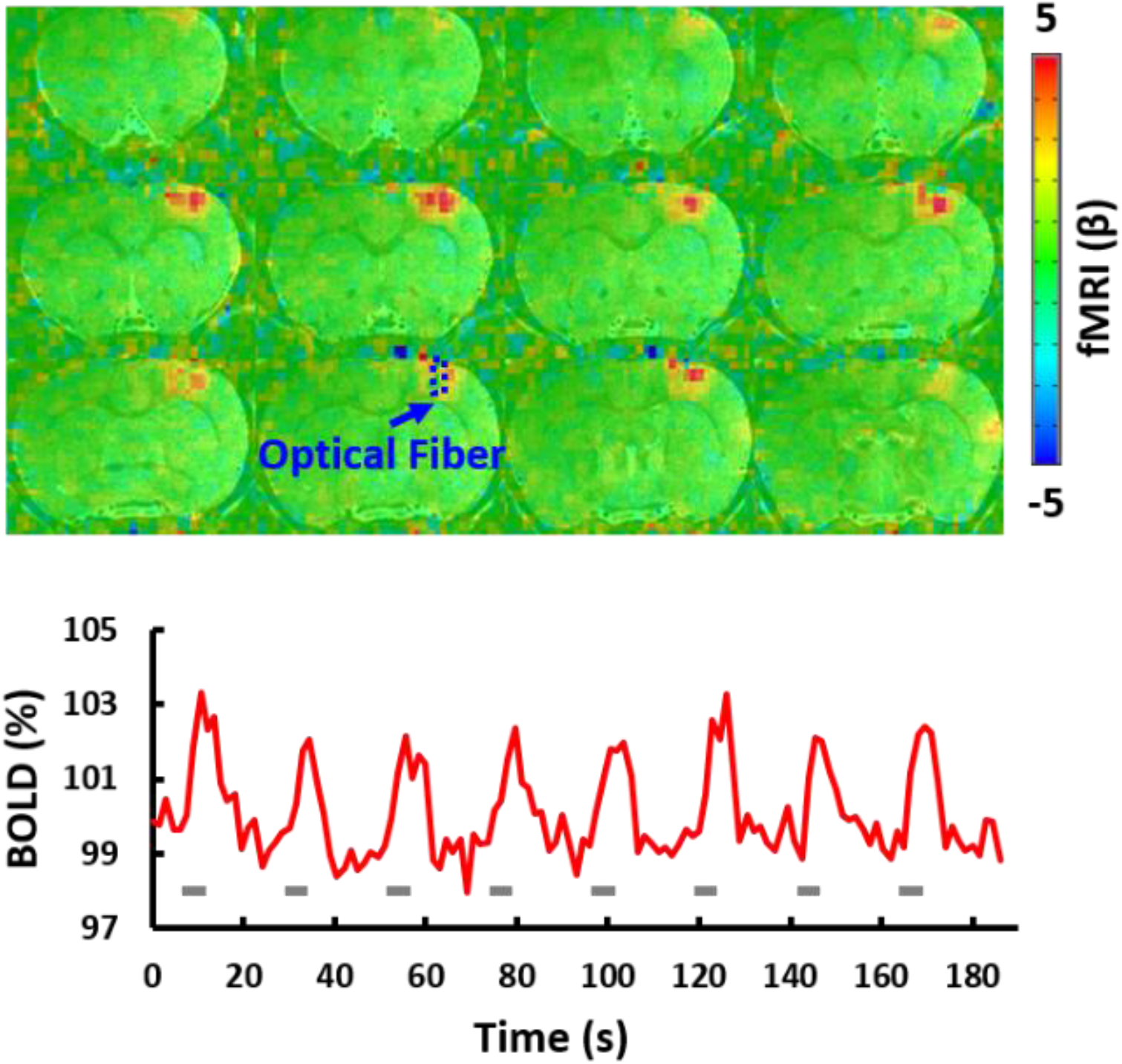
Whole brain BOLD mapping (**a**) and time course (**b**) upon optogenetic stimulation, the blue arrow indicates the optical fiber. Grey bars indicate the optogenetic stimulation.

